# CaMKIIα and CaMKIIβ hub domains adopt distinct oligomeric states and stabilities

**DOI:** 10.1101/2023.10.09.561517

**Authors:** Can Özden, Sara MacManus, Ruth Adafia, Alfred Samkutty, Ana Torres Ocampo, Scott C. Garman, Margaret M. Stratton

## Abstract

Ca^2+^/calmodulin-dependent protein kinase II (CaMKII) is a multidomain serine/threonine kinase that plays important roles in the brain, heart, muscle tissue, and eggs/sperm. The N-terminal kinase and regulatory domain is connected by a flexible linker to the C-terminal hub domain. The hub domain drives the oligomeric organization of CaMKII, assembling the kinase domains into high local concentration. Previous structural studies have shown multiple stoichiometries of the holoenzyme as well as the hub domain alone. Here we report a comprehensive study of the hub domain stoichiometry and stability in solution. We solved two crystal structures of the CaMKIIβ hub domain that show 14mer (3.1 Å) and 16mer (3.4 Å) assemblies. Both crystal structures were determined from crystals grown in the same drop, which suggests that CaMKII oligomers with different stoichiometries likely coexist. To further interrogate hub stability, we employed mass photometry and temperature denaturation studies of CaMKIIβ and α hubs, which highlight major differences between these highly similar domains. We created a dimeric CaMKIIβ hub unit using rational mutagenesis, which is significantly less stable than the oligomer. Both hub domains populate an intermediate during unfolding. We found that multiple CaMKIIβ hub stoichiometries are present in solution and that larger oligomers are more stable. CaMKIIα had a narrower distribution of molecular weight and was distinctly more stable than CaMKIIβ.

## INTRODUCTION

Ca^2+^/calmodulin-dependent protein kinase II (CaMKII) is a serine/threonine kinase that is crucial to memory formation, cardiac pacemaking, and fertilization. The unique multimeric structure of CaMKII is driven by the C-terminal hub domain. The N-terminal kinase domain is tethered to the hub domain through a regulatory segment and flexible linker. The hub domains assemble as two stacked rings, which form the core of the CaMKII holoenzyme.^1,2^ CaMKII is highly conserved across species, with the exception of the linker region, and is expressed by four paralogous genes in humans (α, β, δ, and γ).^3,4^ CaMKII is a metazoan protein, and homologs of full-length CaMKII have been found in choanoflagellates.^5^ Homologs of the CaMKII hub domain have also been found in bacteria and plants.^6^

Structural studies have shown that each ring of the hub domain has 6-8 subunits, forming oligomers with 12-16 total subunits.^5-11^ To date, the human and mouse CaMKII hub domains of α, δ, and γ have been crystallized as either 12mer or 14mer complexes.^5,7,10^ A CaMKII hub-like protein from *Chlamydomonas reinhardtii* has been crystallized as an 18mer complex, which indicates the possibility of higher stoichiometries.^6^ Upon comparative analysis, it has been proposed that different stoichiometries are facilitated by the changes in beta sheet curvature and/or additional intrachain hydrogen bonds.^6^ A recent study revealed a putative model of the first 16mer complex of a CaMKIIβ holoenzyme using negative stain electron microscopy (EM).^12^ The same study showed that both CaMKIIα and CaMKIIβ holoenzymes form mostly 12mers along with a lower population of 14mers.

Mass photometry (MP) measures the molecular weight of single particles in solution.^13,14^ This technique has been used to show that CaMKIIα holoenzymes in solution cluster around a molecular weight of a 10-12mer,^15,16^ whereas the CaMKIIα hub domains form larger species, around 16mers.^16^ The CaMKIIα hub domain is significantly more stable than the CaMKIIα holoenzyme in terms of temperature stability and oligomerization.^16,17^ Another study measured the melting temperature of the monomeric *C. elegans* CaMKII variant to be roughly 63 °C.^18^ Overall, the stability of CaMKII variants has been largely understudied.

Here, we report two human CaMKIIβ hub domain crystal structures at <3.5 Å with 14- and 16mer stoichiometries. We use MP and circular dichroism (CD) to interrogate the stoichiometry and stability in solution of the CaMKIIβ hub domain, and we compared this to the CaMKIIα hub. It has been previously shown that the hub domain impacts CaMKII activity,^19^ so it is important to consider distinctions between variants in understanding how stoichiometry and hub dynamics impact CaMKII regulation.

## RESULTS

### Two CaMKIIβ hub stoichiometries from a single crystallization drop

Initial CaMKIIβ hub crystals were canoe shaped (Figure 1a). A 14mer hub structure was determined at 3.1 Å resolution using a data set collected from a crystal on the right side of the drop (Figure 1a, b, PDB: 7URW). The space group (C222_1_) and unit cell dimensions (104.24, 183.19, 108.37) of this structure are comparable to a recently published CaMKIIα hub 14mer (PDB: 7REC).^12^ Crystal packing and oligomer dimensions (112 Å x 112 Å x 60 Å, Figure 1b, c, Table 1) of both structures are nearly identical.

**Figure 1.**
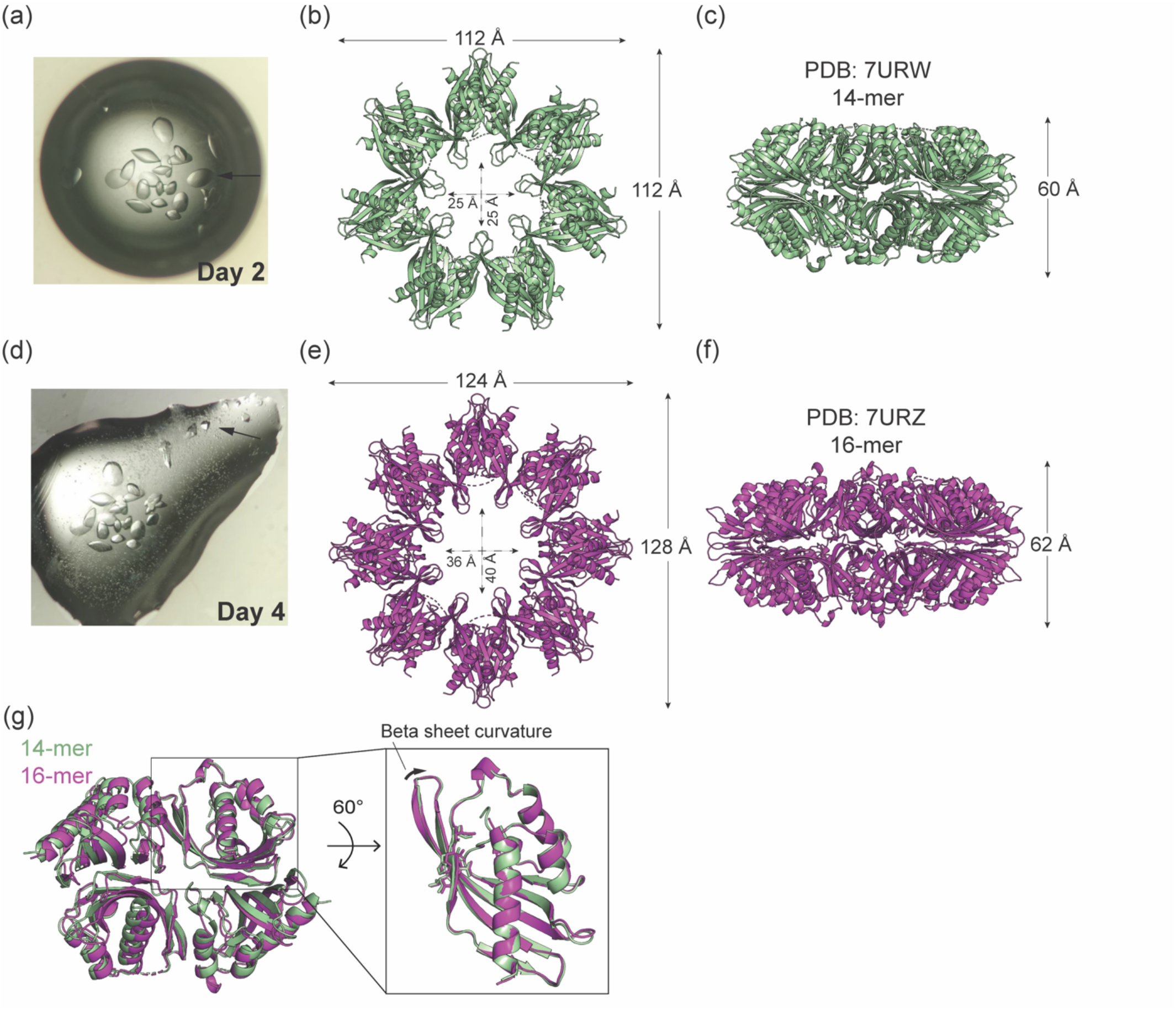
Two CaMKIIβ hub domain stoichiometries from one crystallization drop. (a) Protein crystals shown at the two-day timepoint. The arrow indicates the crystal used for data collection of crystal structure presented in (b). 14mer hub domain crystal structure (PDB code: 7URW) shown (b) top-view and (c) side-view. (d) Protein crystals with different morphology appeared in the same drop at the four-day timepoint. The arrow indicates the crystal used for data collection of crystal structure presented in (e). 16mer CaMKIIβ hub domain crystal structure (PDB code: 7URZ) shown (e) top-view, (f) side-view. (g) Overlay of tetramers from 14mer (green, PDB: 7URW) and 16mer (magenta, PDB: 7URZ) structures. On the left, a tetramer of the hub domains is shown. Structures are aligned along the core α-helix and the first β-sheet of the domain in the box. The inset shows an orientation of the subunit where the curvature is highlighted, as evidenced by the offset of residues highlighted as sticks for clarity. Increased beta sheet curvature facilitates larger ring formation.

Two days later, crystals with a different morphology (tetragonal bipyramid with 4-fold symmetry), as well as many microcrystals, appeared in the same drop (Figure 1d). From these crystals, we determined a 16mer structure at 3.4 Å resolution (PDB: 7URZ). There are 4 hub domains in the asymmetric unit cell with a P4 symmetry. The space group (P4_2_2_1_2) and unit cell dimensions (81.34, 81.34, 180.04) are distinct from the tetradecameric structure. Comparing this 16mer hub structure to the hub of the 16mer holoenzyme EM structure^11^, the outer diameter measurements are comparable (128 Å x 125 Å x 62 Å, Figure 1e,f, Table 1).

In the structures presented here, the curvature of the central β-sheets changes with stoichiometry, as described previously (Figure 1g).^6^ The difference in angle between β-sheets in CaMKIIα 12mer compared to a 14mer is ∼8.6°.^5^ The difference in angle is closer to 3° when we compared the 14mer and 16mer CaMKIIβ hubs. We would expect that as ring size increases, the angle between subunits will not change as drastically. The 16mer is subtly larger in all dimensions compared to 14mer (Figure 1c, f). Comparison of the inside cavities shows that the 16mer structure has an elliptical cavity (36 Å x 40 Å) whereas the 14mer structure cavity is circular (25 Å x 25 Å, Figure 1b, e).

The C-terminus of each hub domain extends from the final β-sheet (β6), which faces the central pore of the donut (Figure S1a). The construct used for the CaMKIIβ hub structures presented here includes all C-terminal residues, but the last 3-5 residues are only well-resolved in 1 out of 11 unique chains between the two structures. We compared an additional 14mer CaMKIIβ hub structure deposited at higher resolution (2.6 Å, PDB code: 7URY)^20^, where the last 3-5 residues are resolved in 5 out of 7 unique chains. We observe that these terminal residues fold back toward the central pocket of their own hub domain (Figure S1a). Two proline residues facilitate turns at the C-terminus (Figure S1b, c).

### CaMKIIβ hub stability

Since multiple stoichiometries can be formed in crystals, we sought to interrogate stoichiometry and its effects on stability in solution. We performed differential scanning calorimetry (DSC) on the CaMKIIβ hub, which showed an apparent melting temperature (T_m_) of 102.2 °C (Figure S2a). We then used circular dichroism (CD) to measure protein unfolding, and, as expected from the DSC data, a standard temperature melt to 100 °C did not completely unfold the protein (Figure S2b). We performed the temperature melt in the presence of 2M guanidine hydrochloride (GdnHCl) and observed two distinct, cooperative unfolding transitions (Figure 2a). This process was irreversible, so all values reported here are phenomenological (Figure S3a, b). The first transition (N ⇌I) had a T_m_ of 56.5 °C and an m-value of 0.18 ± 0.04 cal mol^-1^ °C^-1^. The second transition (I ⇌U) had a T_m_ of 81.8 °C and an m-value of 0.35 ± 0.06 cal mol^-1^ °C^-1^ (Table 2). We also performed guanidine melts on the CaMKIIβ hub (Figure 2b). The oligomer is fully unfolded at 4.5 M GdnHCl and evidently transitions through an intermediate, but these data were not easily fitted to a 2- or 3-state equation.

**Table 2:** Thermodynamic parameters for N ⇌I and I ⇌U transitions. Mean and standard deviation for the calculated slope, ΔG_app_, and T_m_ for the two transitions observed in the temperature melt; n=9.

| | slope (kcal mol<sup>-1</sup> K<sup>-1</sup>) | $\Delta G_{app}$ (kcal mol<sup>-1</sup>) | $T_m$ (°C) |
| --- | --- | --- | --- |
| $N \rightleftharpoons I$ | $0.18 \pm 0.04$ | $10 \pm 2$ | $56.5 \pm 1.9$ |
| $I \rightleftharpoons U$ | $0.35 \pm 0.06$ | $28 \pm 5$ | $81.8 \pm 1.4$ |

**Figure 2.**
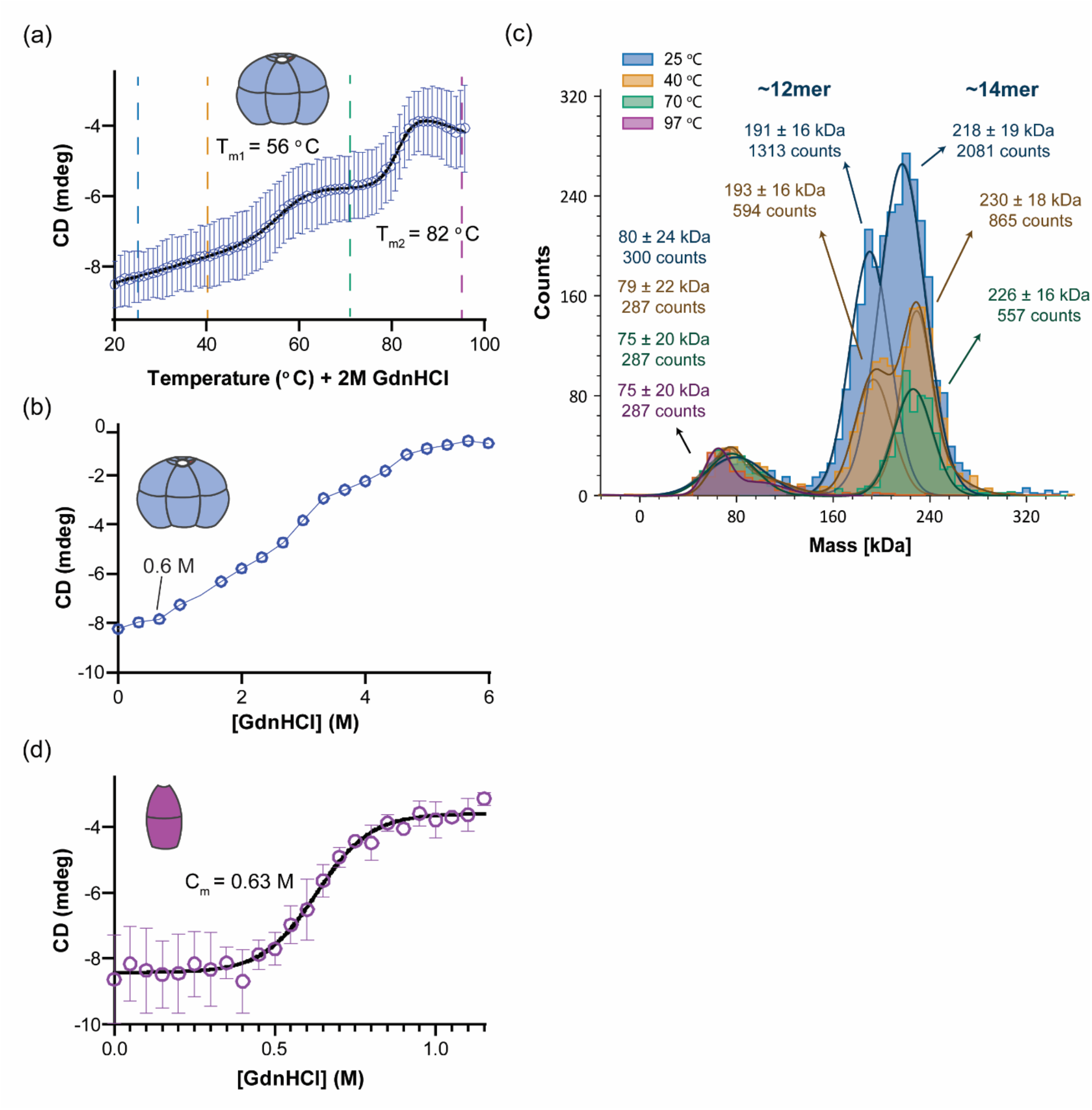
CaMKIIβ hub unfolding. (a) Temperature melt of CaMKIIβ hub oligomer in the presence of 2M GdnHCl. Data points represent average CD signal at 218 nm (n=9). Data are fit to a 3-state non-linear regression. Vertical dashed lines indicate temperatures at which MP measurements were performed (2c). (b) Guanidine melt of CaMKIIβ hub oligomer. Blue data points represent average CD signal at 218 nm (n=9). (c) Representative MP histogram for CaMKIIβ hub samples (n = 8) that have been incubated in 2M GdnHCl and heated to the indicated temperature. Each count indicates a single molecule. (d) Guanidine melt of CaMKIIβ hub dimer mutant (F585A + L622M). Data points represent average CD signal at 218 nm (n=9). Data are fit to a 2-state non-linear regression.

### Defining the unfolding mechanism

We utilized mass photometry (MP) to determine the oligomeric state of the CaMKIIβ hub during the unfolding process. We performed MP measurements of samples collected along the unfolding trajectory as shown in Figure 2a, sampling temperatures representing each state along the unfolding pathway in the presence of 2 M GdnHCl (25 °C, 40 °C, 70 °C, and 97 °C) (Figure 2c, S4a-c). Since MP is performed at significantly lower concentrations, we verified that a dilution series maintains the same populations with and without GdnHCl (Figure S5a, b). At all temperatures, there is a small population around 75 kDa. Control measurements indicated that this is partially contributed by guanidine, and likely partially contributed by lower-order species or unfolded monomers (Figure S4d). Before the first transition (25 °C and 40 °C), there are two distinct CaMKIIβ hub populations that represent ∼12mers and ∼14mers. The lower mass peaks are centered around 191 ± 16.5 kDa (25 °C) and 193 ± 15.6 kDa (40 °C), which correspond to 12mers (180 kDa) along with a small population of 14mers (210 kDa). The higher mass peaks are centered around 218 ± 19.2 kDa (25 °C) and 230 ± 17.7 kDa (40 °C), which correspond to a mixture of 14mers (210 kDa) and 16mers (240 kDa). Notably, there is a population shift from 218 kDa to 230 kDa between 25 °C and 40 °C, indicating that the larger species are selected for at higher temperatures. After the first unfolding transition (70 °C), the lower-mass oligomeric peak disappears and a single peak with a mass of 226 ± 16.0 kDa is observed, representing a mixture of 14mers and 16mers. After the second unfolding transition (97 °C), no oligomeric peaks remain. We cannot measure whether the proteins are all monomeric at 97 °C (the resolution of the Refeyn One MP cannot detect individual hub monomers of 15 kDa), but it is likely that all hub domains have fully unfolded because the CD shows all secondary structure was melted at this temperature (Figure 2a, S3b).

Since MP can distinguish larger species but not smaller ones, we wanted to use a different technique to assess the stability of a smaller hub unit. There is mounting evidence that upon dissociation, CaMKII releases vertical dimer units.^21,22^ We designed a double-mutant (F585A + L622M) that disrupts the lateral interfaces in the CaMKIIβ hub domain to form predominantly vertical dimers. A single mutation has not sufficed to eliminate all oligomerization in solution.^5,22^ Size-exclusion chromatography paired with multi-angle light scattering confirmed that the double-mutant we created (herein referred to as dimer) migrated as a single peak and had a calculated mass of around 36 kDa, whereas the WT hub domain had two mass peaks: 207 kDa and 176 kDa (Figure S6a). This confirmed our double-mutant forms only dimers in solution. Additionally, the CD signatures of the dimer and the WT hub overlap well, indicating that these mutations do not affect monomer folding (Figure S6b). We performed a guanidine melt on the dimer, which gave a C_m_ of 0.63M GdnHCl (Figure 2d). We directly compared the unfolding of the CaMKIIβ hub oligomer to that of the hub dimer and observed that the oligomer starts to unfold around 90 °C whereas the dimer is completely unfolded by 60 °C (T_m_ = 51.7) (Figure S2b, c).

### CaMKIIα hub displays differences in stability compared to CaMKIIβ hub

CaMKIIα and CaMKIIβ hub domains share 77% sequence identity – but do they act similarly in solution? In MP measurements, the CaMKIIα hub formed a single population centered at a 16mer size (240 kDa), as opposed to the two populations observed for the CaMKIIβ hub (Figure S7, 8). We performed temperature denaturation melts on the CaMKIIα hub in the presence of 2M GdnHCl. These showed two unfolding transitions, with T_m_ values of 71.8 °C and 85.2 °C (Figure 3a), both higher than those of the CaMKIIβ hub unfolding (see Figure 2a). We performed MP on the CaMKIIα hub samples at temperatures before and after each transition (Figure 3b). The single population of the CaMKIIα hub decreased in number across each transition (from 60 °C to 90 °C), indicating oligomer dissociation and unfolding. However, there is no clear change in stoichiometry of the oligomer population across transitions nor an emergence of new populations that would directly account for the two separate transitions.

**Figure 3.**
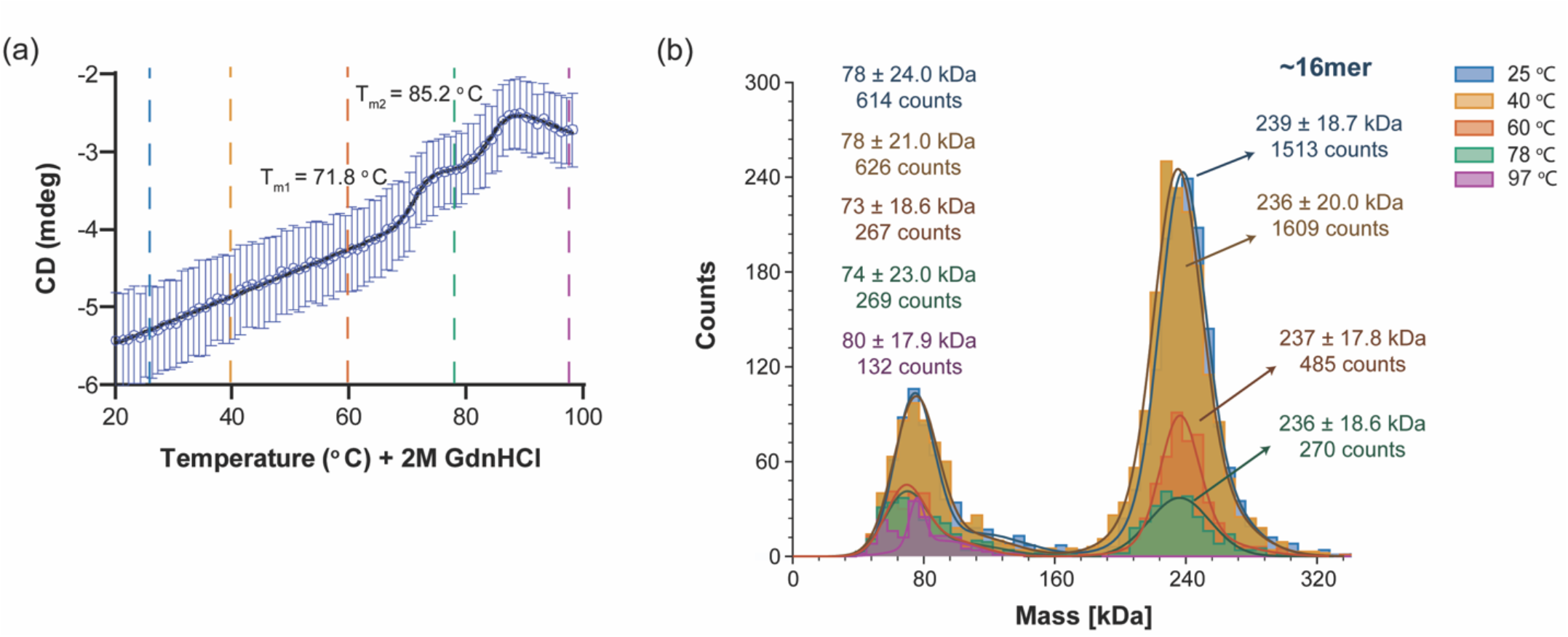
CaMKIIα hub unfolding. (a) Temperature melt of CaMKIIα hub oligomer with 2M GdnHCl. Blue data points represent average CD signal at 218 nm (n=6). Data are fit to a 3-state non-linear regression. Vertical dashed lines indicate temperatures at which MP measurements were performed (2b). (b) Representative MP histogram for CaMKIIα hub samples (n = 6) that have been incubated in 2M GdnHCl and heated to the indicated temperature. Each count indicates a single molecule. Note that the 25 °C population (blue) is overlapping with the 40 °C population (yellow).

## DISCUSSION

The CaMKII holoenzyme is assembled by the hub domain, which oligomerizes to form a donut-like structure. The N-terminus of the hub domain forms a helix which connects to the catalytic domain in the full-length holoenzyme on the outside of the donut (Figure S1a). If the C-terminus was extended, it is unclear where these residues would travel to – this is an important factor to consider with fusion proteins and tags added at the C-terminal end of CaMKII. Of note, CaMKIIδ has an extended C-terminus, which will likely have distinct effects on hub dynamics and stability. Each hub domain harbors a central pocket, which is lined with arginine residues. A previous study determined a structure of the CaMKIIα hub domain bound to an analog of γ-hydroxybutyrate (5-HDC) in this arginine pocket and showed that this compound bound specifically to CaMKIIα.^17^ Other hub structures also trap molecules in this pocket, for example the other 14mer CaMKIIβ hub structure (PDB code: 7URY) shows malic acid bound in the pocket.^20^ The specificity for 5-HDC is striking, and rather remarkable given the high sequence conservation between CaMKII variants. A major difference between the hub domains is that in CaMKIIα there are three arginines in the pocket, while in CaMKIIβ there are only two, and the third is substituted with cysteine (Figure S1b, c). Another difference is the sequence of the five C-terminal residues (Figure S1c), where both segments would be flexible but the sequences are different. As noted, our constructs have full-length C-termini compared to other CaMKIIα structures which are terminated early; this may also affect stoichiometry and it will be important to interrogate this further.

We were intrigued to capture two distinct oligomeric states grown within the same crystallization drop, suggesting that these species coexist and/or interconvert. While the biological implications of stoichiometry are not clear, it is possible that these allowable conformations lend CaMKII its unique properties such as subunit exchange.^5,21^ A recent study has shown that CaMKIIβ has different activation properties compared to CaMKIIα.^19^ CaMKIIα requires higher concentrations of Ca^2+^/calmodulin for activation, whereas CaMKIIβ was active at much lower concentrations. This difference may be attributed to different stoichiometric states.

Both CaMKIIα and β hub domains are extremely stable, requiring temperatures of over 90 °C for unfolding.^16^ In fact, in order to observe unfolding in an achievable temperature range, we needed to add denaturant (2 M GdnHCl). Surprisingly, both hub domains unfolded in a 3-state reaction, but these reactions are distinct from one another despite the high sequence similarity between these domains (Figure S8a). CaMKIIβ hub undergoes its first unfolding transition at a lower temperature compared to CaMKIIα. From MP measurements, we interpret this first transition as the 12mer oligomers unfolding, while the 14/16mer oligomers remain (Figure S8a-e). The second transition is the remaining 14/16mer oligomers dissociating and unfolding. We interpret this to mean that the 14/16mer population is more stable; the 14/16mer has more buried interfaces, suggesting that this stability can be attributed to favorable enthalpic contributions provided by the higher oligomeric state. The CaMKIIα unfolding mechanism appears to be more nuanced, as the oligomeric state (16mer) does not obviously change during unfolding. A possible explanation for the first unfolding transition (71 °C) is that the oligomeric ring cracks at one vertical interface, creating a spiraled ring which has been previously reported.^5^ The second transition would then be all spiraled oligomers dissociating and unfolding. It also notable that the CaMKIIα transitions (closed to open ring/dissociation and unfolding) are closer energetically (Δ13 °C) compared to the transitions for CaMKIIβ (Δ26 °C). The second transition for CaMKIIα and CaMKIIβ hubs is overlapping (Figure S8a), it is possible that full dissociation and unfolding is the same energetically for both variants.

The molar ellipticity of the hub dimer we created is the same as that of the oligomer, so any change in CD signal observed during the denaturation is likely attributed to monomer unfolding rather than disassembly into dimers. Both hub domains displayed a large native baseline slope over the first 40 – 60 degrees of the experiment. We interpret this slope as oligomers (regardless of stoichiometry) disassembling into dimers and unfolding immediately (Figure 4a), which is reflected in the MP measurements where the oligomeric peak(s) counts decrease steadily over this temperature range (Figure 2c, 3b). We cannot rule out precipitation occurring during the measurement; however we do not observe any large species in MP. We were unable to fit the GdnHCl melt of the CaMKIIβ hub domain to a three-state curve, as the intermediate was not well-sampled under these conditions. However, the native baseline only begins to slope after ∼0.5 M GdnHCl. This corresponds with the C_m_ of the CaMKIIβ hub dimer (∼0.6 M), indicating that dimers are unfolding at this point, likely because of oligomer dissociation. There are two visible transitions, one around 3 M GdnHCl and a second around 4.5 M GdnHCl. These transitions likely represent the same processes we visualized in the temperature + GdnHCl denaturation melts.

**Figure 4.**
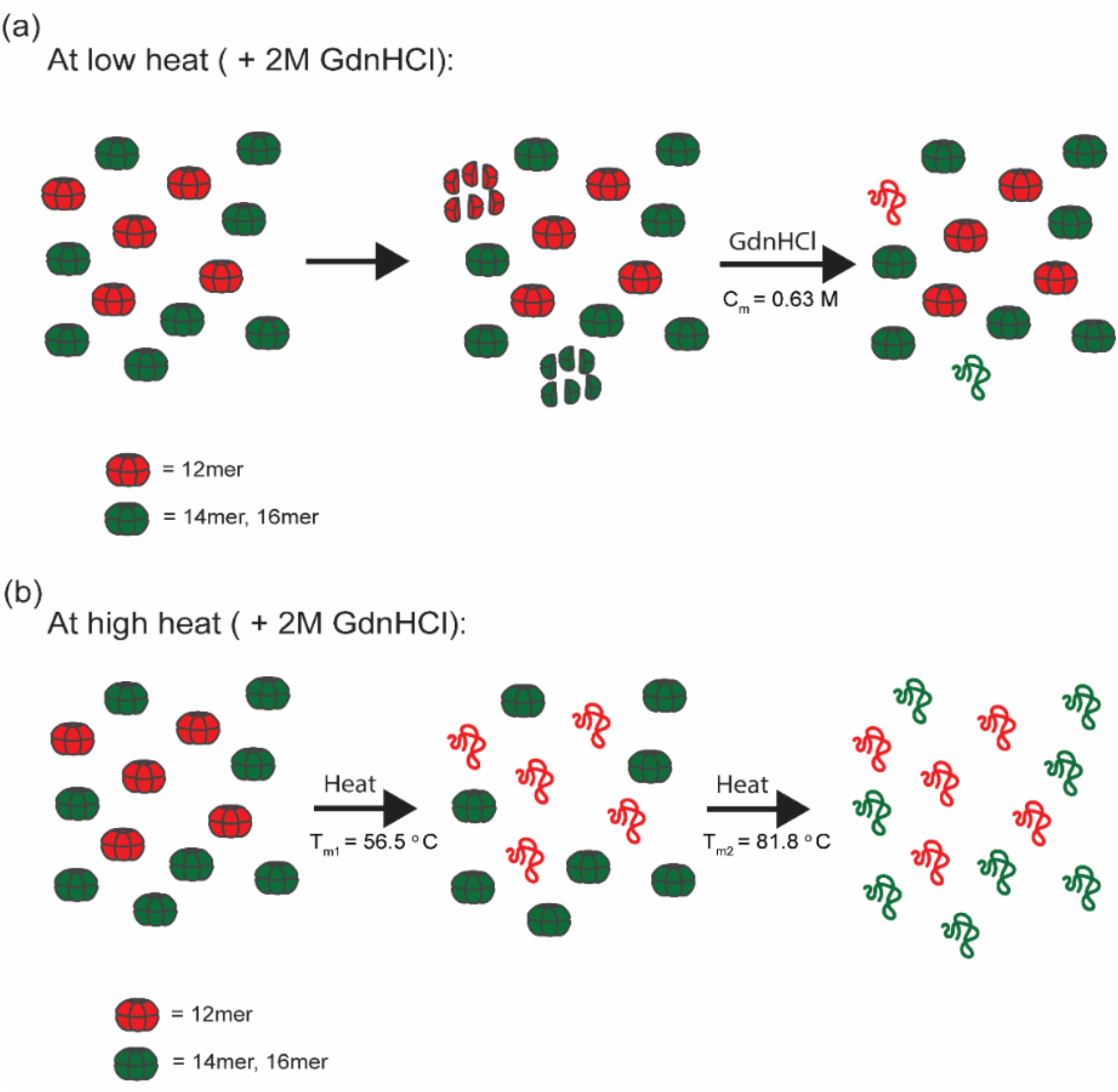
Schema of proposed mechanism of unfolding for CaMKIIβ hub oligomer during temperature melts in the presence of 2 M GdnHCl. (a) At low heat, some oligomers disassemble into dimers regardless of stoichiometry. The dimers immediately unfold. (b) At high heat, 12mers first disassembly and unfold in a cooperative transition. 14mers and 16mers disassemble and unfold in a second cooperative transition.

It is clear from both crystallography and electron microscopy studies that CaMKII holoenzymes and hub domains can adopt different stoichiometries. This adds to the complexity of this enzyme, which not only can form heterooligomeric complexes, but can also have mixed oligomeric states of heterooligomers. The physiological relevance of these differences remains unclear but is important to interrogate moving forward. It is possible that the allowance of multiple stoichiometries facilitates subunit exchange between CaMKII holoenzymes.^5,21^ Our study has provided clear evidence that, despite nearly identical sequences, CaMKIIα and β hub domains adopt different stoichiometries in solution and have distinct stabilities (Figure S8). This provides a framework for assessing the differences observed in activation and regulation between CaMKIIα and β holoenzymes and how they may be related to dynamics and stability. Before addressing such questions, it would be necessary to determine whether full-length CaMKII holoenzymes exhibit the same stoichiometries and disassembly properties as those observed here in hub domain holoenzymes.

## Material and Methods

### Protein purification

The CaMKIIβ full-length hub domain (residues 534-666) was expressed in BL21 DE3 cells with a N-terminal hexahistidine tag. The cells were induced at 18°C by adding 1 mM IPTG and grown overnight. The cells were resuspended in Buffer A (25 mM Tris-HCl pH 8.5, 150 mM KCl, 40 mM imidazole, and 10% glycerol, supplemented with commercially available protease inhibitors (0.5 mM benzamidine, 0.2 mM 4-(2-aminoethyl)benzenesulfonyl fluoride hydrochloride (AEBSF), 0.1 mg/mL trypsin inhibitor, 0.005 mM leupeptin, 1 μg/mL pepstatin), 1 μg/mL DNase/50 mM MgCl_2_ were added, and the cells were then lysed. All subsequent steps were performed at 4°C using an ÄKTA pure chromatography system (GE). Cell lysate was loaded onto 5 mL His Trap FF Ni Sepharose column (GE) and eluted with 50% Buffer B (25 mM Tris-HCl pH 8.5, 150 mM KCl, 1 M imidazole, 10% glycerol). The protein was desalted from excess imidazole using a HiPrep 26/10 Desalting column using Buffer C (25 mM Tris-HCl pH 8.8, 150 mM KCl, 1 mM EDTA, 2 mM DTT, and 10% glycerol). The hexahistidine tag was cleaved with Prescission protease overnight at 4°C. Proteins were eluted from HiTrap Q-FF with a step-gradient from 0% to 17% using Buffer BQ (25 mM Tris-HCl pH 8.5, 1 M KCl, 10% glycerol). Eluted proteins were concentrated and further purified using a Superose 6 size exclusion column equilibrated with gel filtration buffer (25 mM Tris-HCl pH 8.0, 150 mM KCl, 1 mM TCEP, 10% glycerol). Fractions (>95% purity) were pooled, concentrated, flash frozen in liquid nitrogen, and stored at -80 °C until needed.

The CaMKIIβ hub dimer variant was designed by modifying the wild type CaMKIIβ hub domain via two mutations: F585A and L622M (residue numbering based on full-length, non-spliced CaMKIIβ). The variant was expressed in BL21 DE3 cells with an N-terminal hexahistidine tag and purified using the same process as the wild type CaMKIIβ hub. During the size exclusion step, a Superdex 75 size exclusion column was used.

### Crystallization, data collection, and structure determination

Protein crystals of 14mer and 16mer CaMKIIβ hub domain were grown in the same drop using hanging drop vapor diffusion at 20 °C. The drops contained 1 μL of the well solution and 1 μL of 1.9 mM hub domain. The well solution was 0.1 M imidazole pH 7, 0.15 DL-Malic acid pH 7 and 22% PEG methyl ether 550. Crystals were frozen directly from the well solution. Diffraction data were collected at a wavelength of 1.5418 Å using a Rigaku MicroMax-007 HF X-ray source, which was coupled to a Rigaku VariMax HF optic system (UMass Amherst). The X-ray data was collected at 100 K. The datasets were integrated, merged scaled using HKL-2000. The structure was solved by molecular replacement with Phaser^23^ using single CaMKIIγ hub domain (PDB: 2UX0)^7^ as the search model. Model building was performed using Coot and the refinement was carried out with Refmac5^24^.

### Guanidine melts

Two solutions of 10 μM CaMKIIβ hub were prepared, containing either 0M GdnHCl or 6 M GdnHCl. All protein dilutions were performed using a modified gel filtration buffer suitable for CD (5 mM Tris-HCl pH 8.0, 150 mM KCl, 1 mM TCEP). A stock 6.5 M GdnHCl solution was used (6.5 M GdnHCl, 5 mM Tris-HCl pH 8.0, 150 mM KCl, 1 mM TCEP) to prepare GdnHCl solutions. The two solutions were titrated together to give an array of 10 μM CaMKIIβ hub solutions with a range of GdnHCl concentration. For the CaMKIIβ hub oligomer, these solutions ranged from 0 M to 6 M GdnHCl at steps of 0.33 M. For the CaMKIIβ hub dimer, these solutions range from 0 M to 1.15 M in 0.05 M steps.

Following the titration, protein samples incubated for 20 minutes (CaMKIIβ hub oligomer) or 50 minutes (CaMKIIβ hub dimer). After equilibration, the extent of unfolding in each solution was measured on a Jasco J1500 CD spectrophotometer using 1 mm cuvettes. This was measured using circular dichroism (CD) signal. For each sample, the average CD at 218 nm was recorded over 30 seconds at 25 °C. Additionally, full spectrum scans (195 nm to 260 nm) were recorded for CaMKIIβ hub oligomer and dimer samples in 0M GdnHCl. The raw CD signal (mdegs) was converted to molar ellipticity (*Θ*) for the full spectrum scans via the following conversion: *Θ* = mdegs * M / (10 * L * C) where M is average molecular weight in g/mol, L is path length of the cell, and C is concentration in g/L.

Using a non-linear regression, data were fitted to a two-state curve (Equation 1) in Graph Pad Prism. The parameter values were used to calculate C_m_ values (C_m_ = m4/m5).

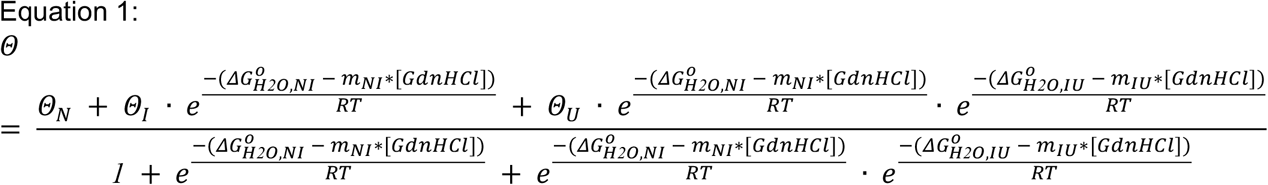

### Temperature melts

Protein samples of 60 μM CaMKIIβ hub and 2 M GdnHCl were prepared at a final volume of 215 μL. Circular dichroism (CD) measurements were performed using a Jasco J1500 CD spectrophotometer. Protein samples in a 1 mm cuvette were heated from 20 °C to 100 °C at a rate of 0.67 °C per minute. Every 1 °C, the CD signal at 218 nm was recorded over 30 seconds. Full spectrum CD scans (205 to 260 nm) were taken every 10 °C. Eight temperature melt replicates were performed. Using a non-linear regression, data were fitted using Equation (2) in GraphPad Prism, where N is the native protein, I is the intermediate, U is the unfolded protein, NI is the first transition, and IU is the second transition. The average m4, m5, m6, and m7 values were calculated over all replicates. These values were used to calculate the melting temperature for each transition (T_m,1_ = m4/m5 and T_m,2_ = m6/m7), and unfolding energies (m4*RT and m6*RT, respectively) (T = 297 K).

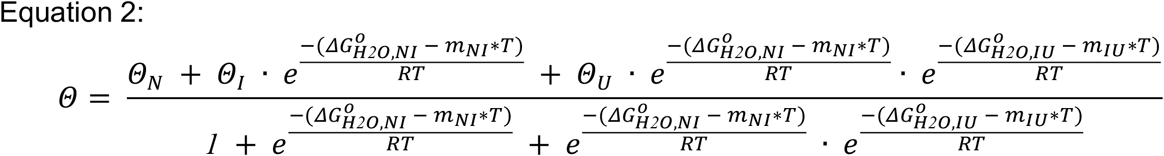

### Mass photometry

All mass photometry measurements were performed on a Refeyn One MP mass photometer. All protein solutions were prepared in a filtered buffer (25mM Tris-HCl pH 8.0, 150mM KCl). Before each experiment, the buffer alone was measured as a control. The buffer was considered clean if the hit count was below 75. For each experiment, a standard mass calibration curve was calculated using TGA at 25 nM (final concentration on the coverslip), apoferritin (440 kDa) at 3.75 nM, and BSA (66.5 kDa) at 15 nM; 5 μL of standard protein was added to 15 μL of MP buffer on a coverslip and the MP was measured for 60 seconds.

For each run, a 200 μL solution of 2 μM protein and 2 M GdnHCl was prepared. The GdnHCl solution was filtered using a 0.22 μm filter before use. The sample was heated to the chosen temperatures using a thermocycler. The sample was held at each temperature for 90 seconds, at the end of which 6 μL were taken and immediately diluted onto the coverslip with 14 μL of MP buffer to perform an MP measurement over 60 seconds. Mass histograms of each run were generated using Refeyn’s DiscoverMP software (n = 6 for CaMKIIa hub, n = 8 for CaMKIIβ hub).

### SEC MALS

Size-Exclusion Chromatography (SEC) coupled with Multi-Angle Light Scattering (MALS) was performed using an Agilent 1260 HPLC system, a DAWN HELEOS II MALS detector, and an Optilab-T-rEX refractive index detector. The SEC column used was a Tosoh TSKGel G3000 SWxl (particle size 5 um, 30 cm length, calibration range 10 kDa – 500 kDa) and was equilibrated with ethanol, distilled water, and finally gel filtration buffer (25 mM Tris-HCl pH 8.0, 150 mM KCl, 1 mM TCEP, 10% glycerol). The SEC-MALS was conducted at a constant flow rate of 0.9 μL/min with an injection volume of 50 μL to ensure accurate separation. Protein samples of CaMKIIβ hub oligomer and dimer were injected at a concentration of 262 mg/mL. The column temperature was maintained at 25 °C throughout the analysis to prevent temperature-induced variations. Data were acquired and processed using ASTRA. The scattering data were analyzed based on the Zimm plot method to determine the molecular weight of the samples. The SEC-MALS system was calibrated using BSA (2 mg/mL) to ensure accurate determination of molecular weights.

## Author Contributions

**Can Özden:** Conceptualization; data curation; formal analysis; investigation; methodology; visualization; writing-original draft; writing-review and editing. **Sara MacManus:** Conceptualization; data curation; formal analysis; investigation; methodology; visualization; writing-original draft; writing-review and editing. **Alfred Samkutty:** Data curation; visualization. **Ana Torres Ocampo**: Data curation; visualization. **Scott Garman:** Formal analysis; visualization. **Margaret Stratton:** Conceptualization; formal analysis; funding acquisition; investigation; methodology; project administration; resources; supervision; visualization; writing-original draft; writing-review and editing.

## Acknowledgements

We thank Stewart Loh and Alejandro Heuck for thoughtful comments on the project. We thank Eddie Esposito for help with DSC experiments. CD and SEC-MALS data were collected at the UMass Amherst Mass Biophysical Characterization Core Facility, RRID:SCR_022357.

## Supplemental Figures

**Figure S1.**
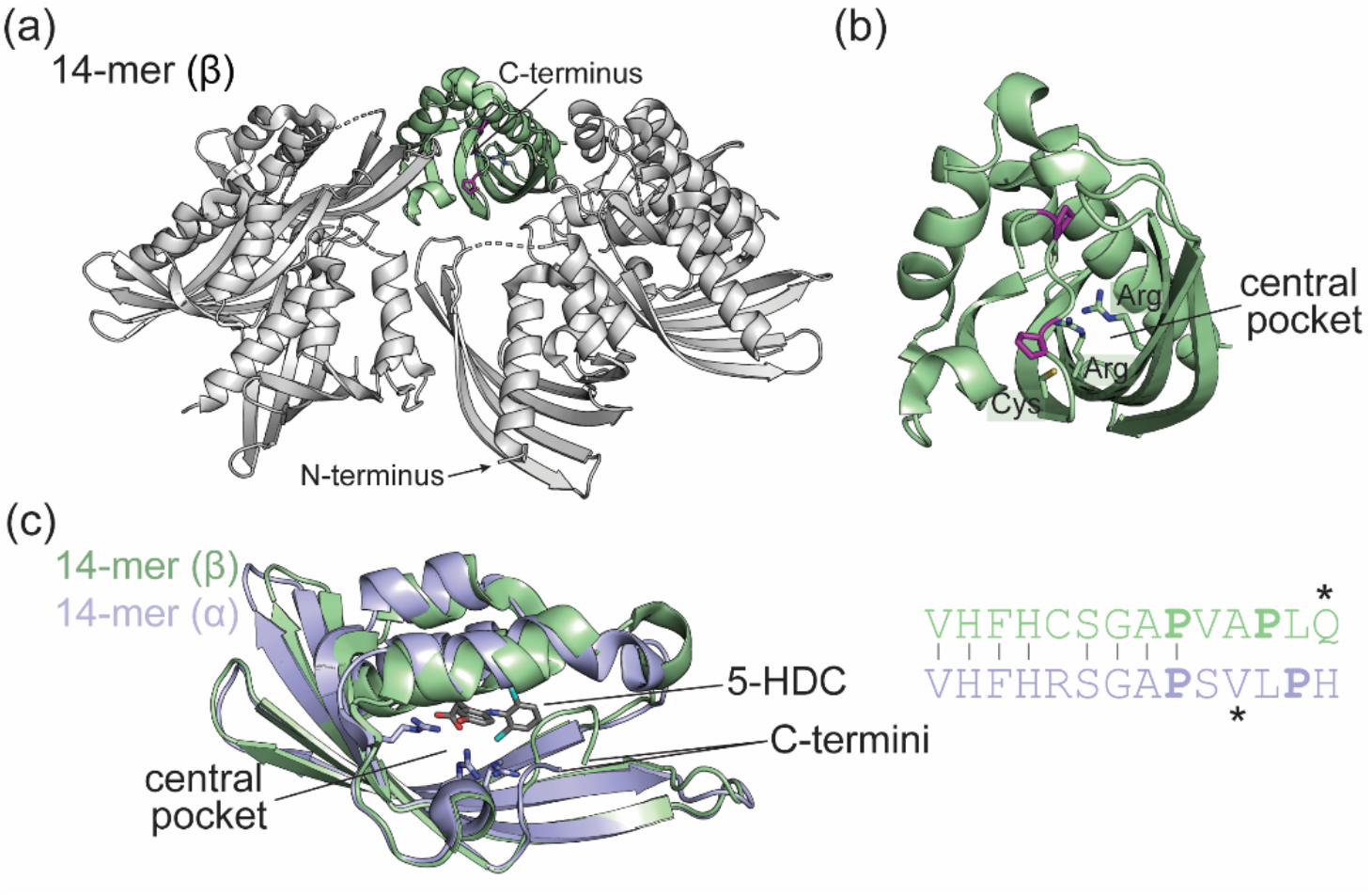
C-terminal residues of CaMKIIβ hub domain extend into the hub pocket. (a) The top ring from the 14mer β hub structure is shown and the hub N- and C-termini are indicated. (b) A zoom-in of the subunit highlighted in green in (a) is shown. The central pocket of CaMKIIβ is lined with two arginine residues (shown as sticks). There is a cysteine (shown as stick) in place of the third arginine found in other CaMKII variants. Two proline residues (highlighted as magenta sticks) cause kinks in the C-terminal tail. (c) The CaMKIIβ hub domain 14-mer that we solved here (green) is overlaid with a CaMKIIα hub domain structure (purple, PDB: 7REC). The central pocket of the CaMKIIα hub is lined with three arginine residues (shown as sticks). There is a molecule bound in the pocket (5-HDC, shown as gray sticks). An alignment of the C-terminal 14 residues of CaMKIIβ (green) and CaMKIIα (light blue) is shown. The asterisks indicate the final residue of the crystallization constructs used – note that the constructs in this manuscript all include the full-length C-terminus.

**Figure S2.**
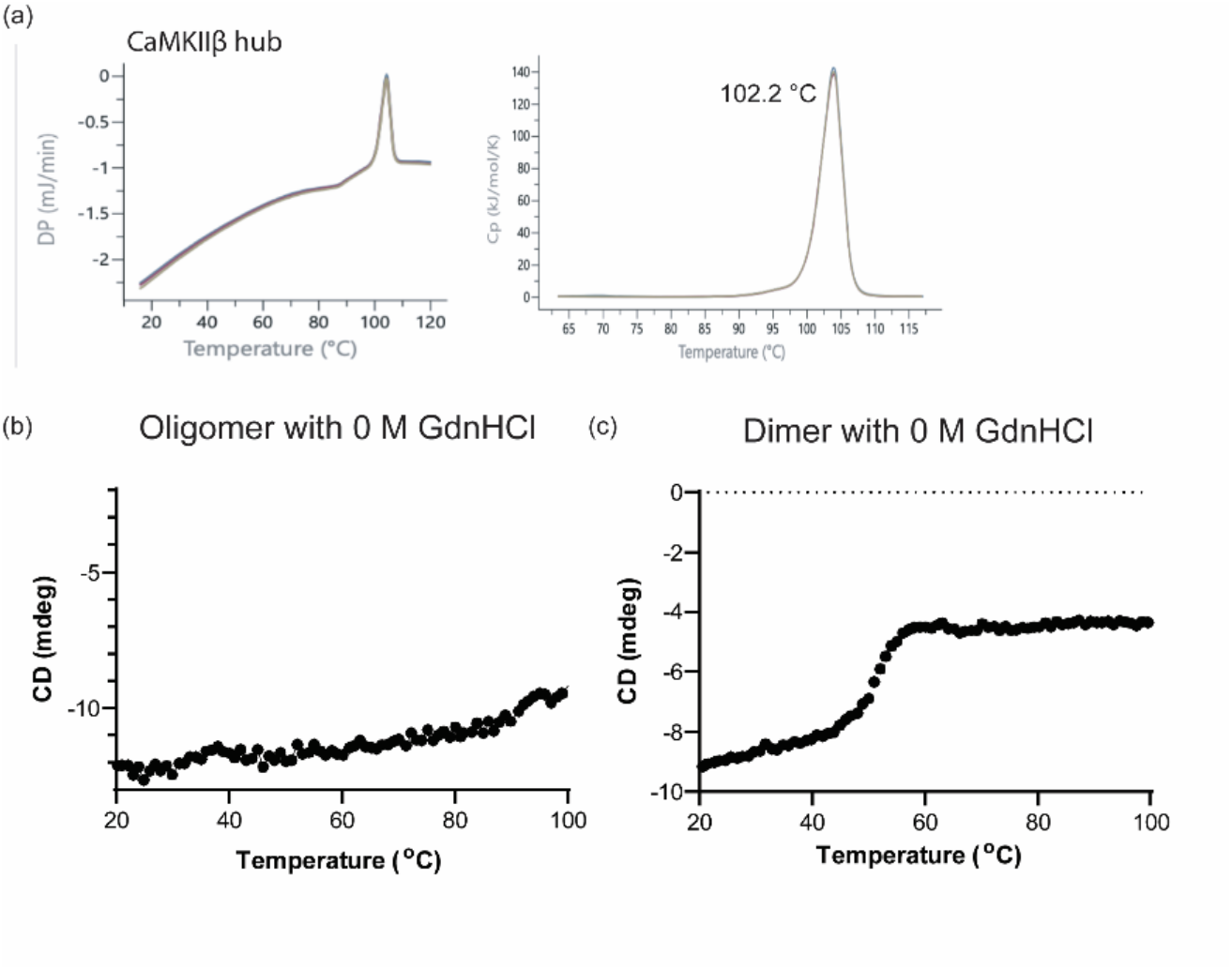
(a) DSC measurements of CaMKIIβ hub oligomer. T_m_ = 102.2 °C. (b) Temperature melt of CaMKIIβ hub oligomer without the addition of GdnHCl. Each data point represents the average CD signal at 218 nm over 30 seconds (n = 1). (c) Temperature melt of CaMKIIβ hub dimer without the addition of GdnHCl. Each data point represents the average CD signal at 218 nm over 30 seconds (n = 1).

**Figure S3.**
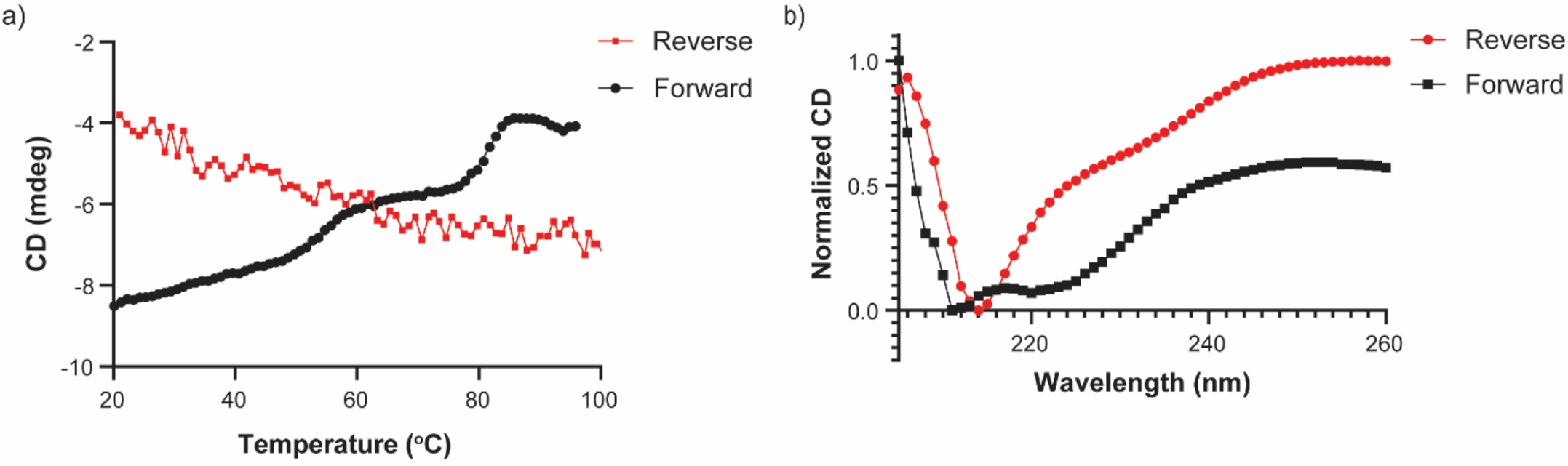
Temperature melts performed on CaMKIIβ hub samples in the presence of 2M GdnHCl are irreversible. (a) Full denaturation curve for both forward and reverse temperature melts. (b) Normalized CD signal across the wavelength spectrum (205-260 nm) of samples at 20 °C. The measurement for the forward temperature melt was performed before heating the sample. The measurement for the reverse temperature melt was performed after cooling the sample from 100 °C to 20 °C.

**Figure S4.**
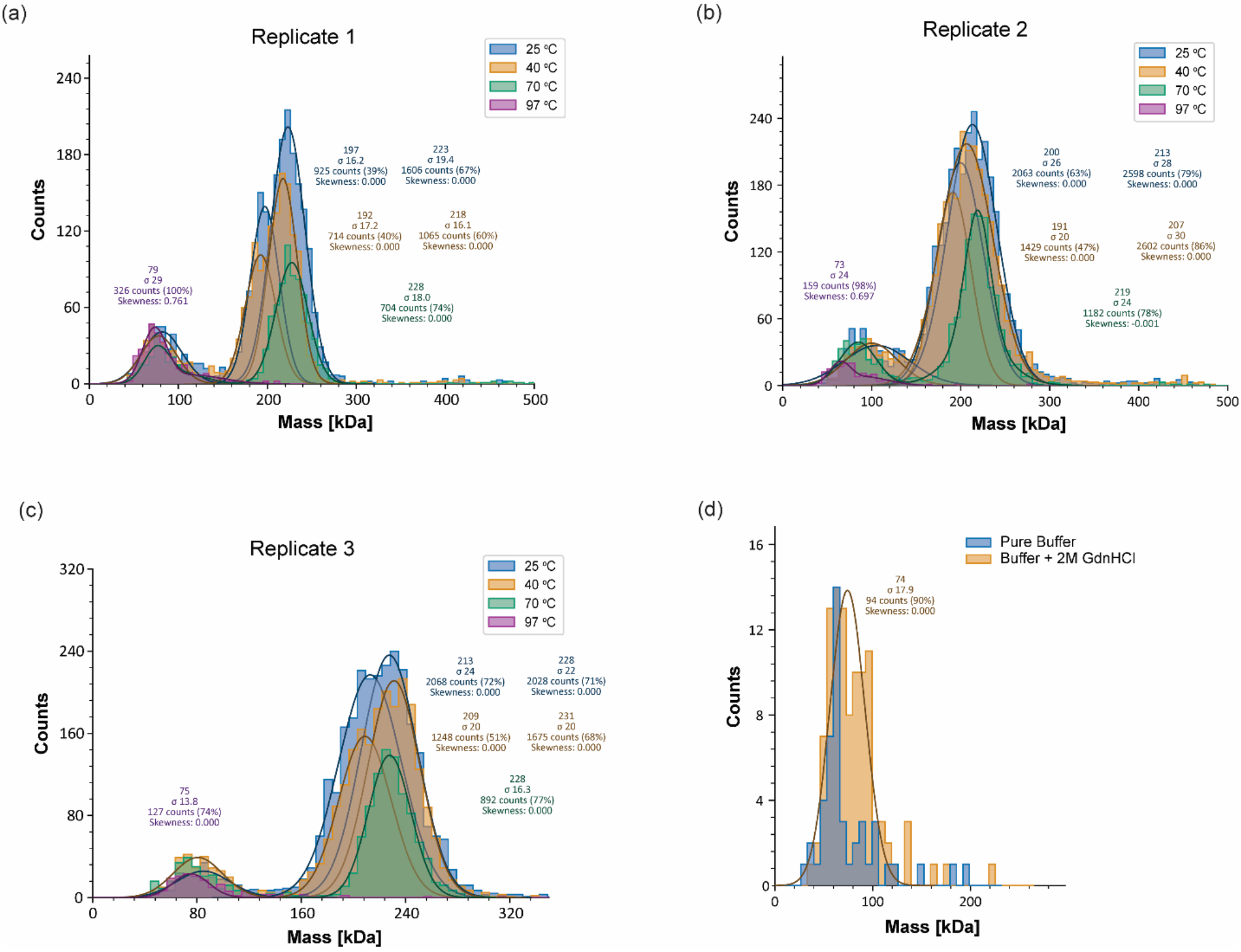
(a-c) Replicates of MP measurements on CaMKIIβ hub samples that have been incubated in 2M GdnHCl and heated to stated temperature. The y-axis on the chart represents the number of counts, each of which indicates an individual recognized molecule. (d) MP measurements comparing buffer and buffer with 2M GdnHCl.

**Figure S5.**
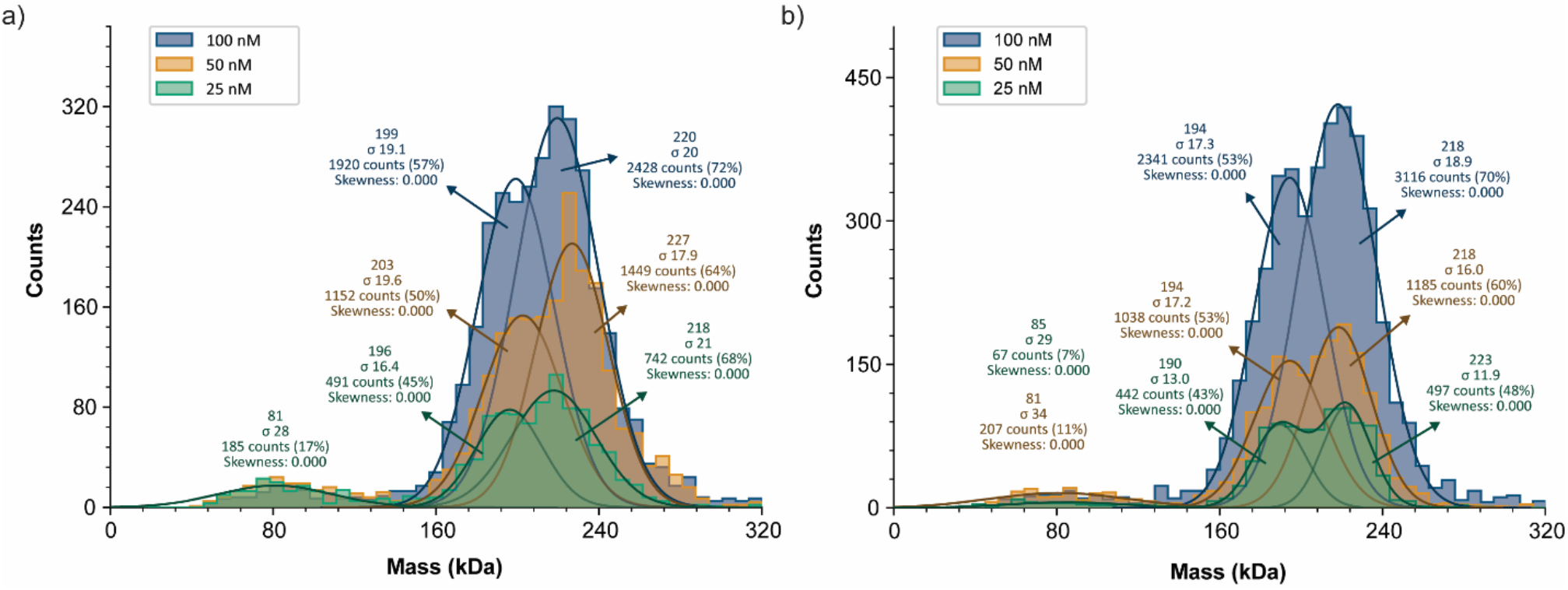
MP measurements on CaMKIIβ hub samples at varying concentrations. (a) MP measurements performed on CaMKIIβ hub samples at the indicated concentrations. (b) MP measurements performed on CaMKIIβ hub samples at the indicated concentrations with 2M GdnHCl.

**Figure S6.**
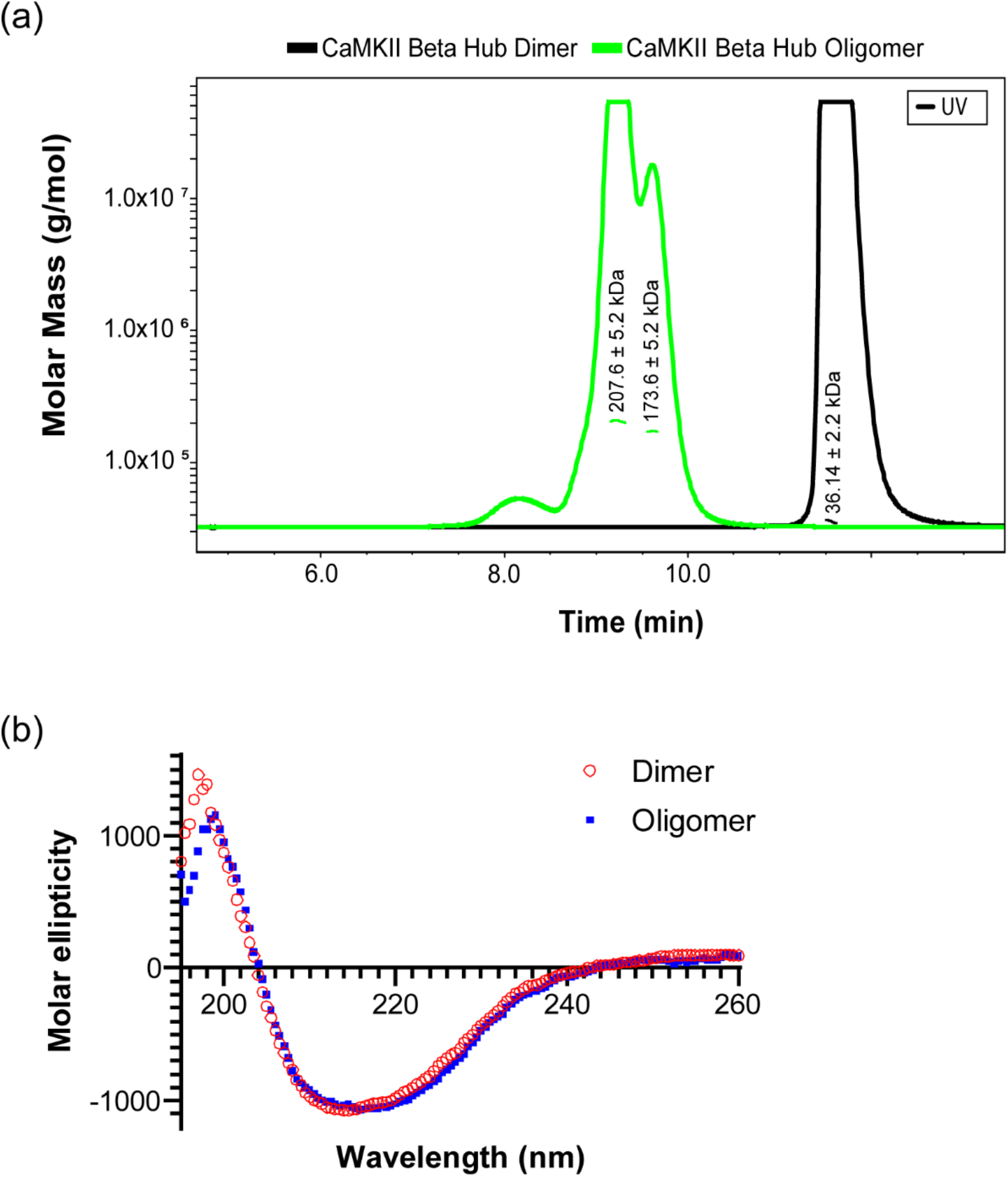
(a) SEC-MALS measurements performed on CaMKIIβ hub oligomer and dimer. The CaMKIIβ hub oligomer produced two populations with molecular masses of 207.6 ± 5.2 kDa and 173.6 ± 5.2 kDa. The CaMKIIβ hub dimer produced one population with a molecular mass of 36.14 ± 2.2 kDa. (b) Molar ellipticity of CaMKIIβ hub oligomer and dimer across the wavelength spectrum (195 nm – 260 nm).

**Figure S7.**
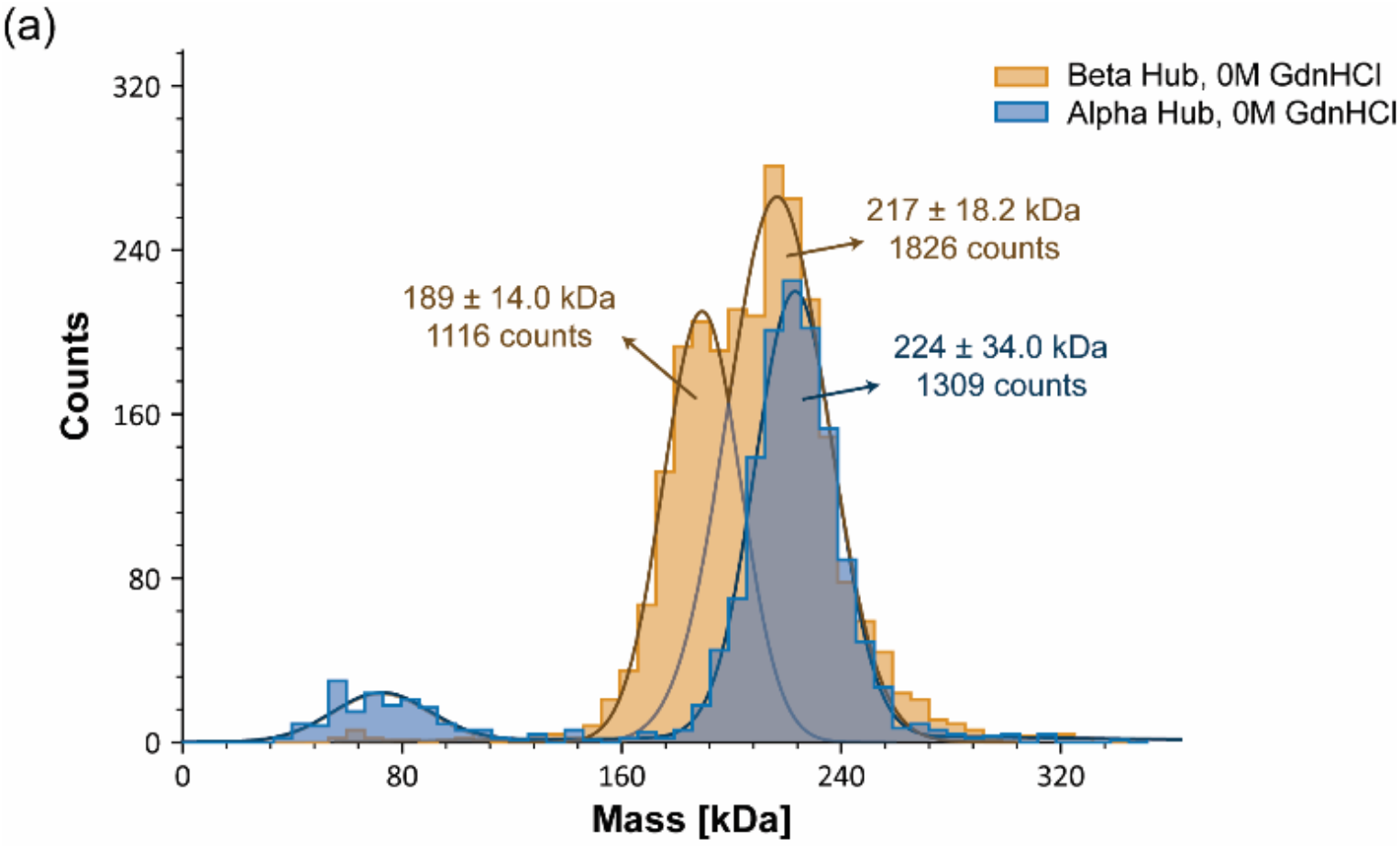
(a) MP measurements performed on CaMKIIβ hub and CaMKIIα hub without the addition of GdnHCl. CaMKIIβ hub exhibits two populations with molecular masses of 189 kDa and 217 kDa. CaMKIIα exhibits a single population at 224 kDa.

**Figure S8.**
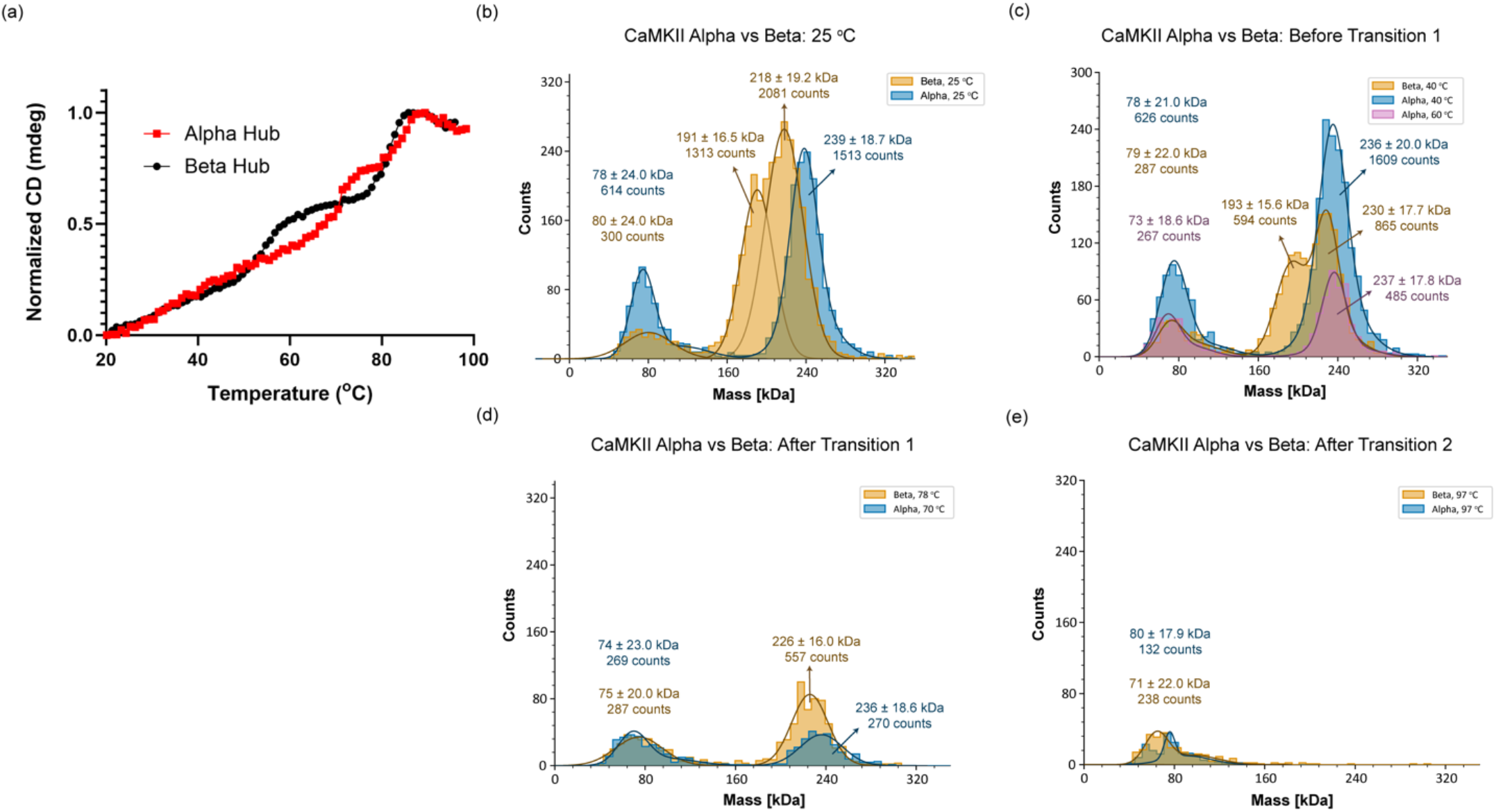
Comparison between CaMKIIβ hub and CaMKIIα hub. (a) Overlay of temperature melts in the presence of 2M GdnHCl. Red data points are CaMKIIα hub, black data points are. CaMKIIβ hub. Fits are excluded for clarity. (b) MP histograms representing CaMKIIβ hub and CaMKIIα hub populations at room temperature. (c) MP histograms representing CaMKIIβ hub and CaMKIIα hub populations before their first unfolding transition. (d) MP histograms representing CaMKIIβ hub and CaMKIIα hub populations after their first unfolding transition and before their second unfolding transition. (e) MP histograms representing CaMδKIIβ hub and CaMKIIα hub populations after their second unfolding transition.

